# Neural dynamics of spontaneous memory recall and future thinking in the continuous flow of thoughts

**DOI:** 10.1101/2024.10.02.616300

**Authors:** Haowen Su, Xian Li, Savannah Born, Christopher J. Honey, Janice Chen, Hongmi Lee

**Affiliations:** Department of Psychological Sciences, Purdue University, West Lafayette, IN 47907, USA; Department of Psychological and Brain Sciences, Johns Hopkins University, Baltimore, MD 21218, USA; Department of Psychological and Brain Sciences, Washington University in St. Louis, MO 63130, USA

## Abstract

The human brain constantly recalls past experiences and anticipates future events, generating a continuous flow of thoughts. However, the neural mechanisms underlying the natural transitions and trajectories of thoughts during spontaneous memory recall and future thinking remain underexplored. To address this gap, we conducted a functional magnetic resonance imaging study using a think-aloud paradigm, where participants verbalize their uninterrupted stream of thoughts during rest. We found that transitions between thoughts, particularly those involving significant shifts in semantic content, activate the brain’s default and control networks. These neural responses to internally generated thought boundaries produce activation patterns resembling those triggered by external event boundaries. Moreover, interactions within and between these networks shape the overall semantic structure of thought trajectories: stronger functional connectivity within the medial temporal subsystem of the default network predicts greater variability in thoughts, while stronger connectivity between the control and core default networks is associated with reduced variability. Together, our findings highlight how the default and control networks guide the dynamic transitions and structure of naturally arising memory and future thinking.

## INTRODUCTION

The human mind is constantly engaged in recalling the past and predicting the future^1,2^. This creates a continuous stream of thoughts, where semantic memory about the world and oneself, episodic recollections of specific events, and future-oriented simulations are intertwined with information from the current environment^3,4^. Understanding the dynamics of this internally generated thought flow can provide crucial insights into how mental representations are organized in the brain and the neurocognitive processes involved in accessing them. For instance, when people recall memories in a continuous stream, the order and transitions between memories follow underlying semantic and temporal associations; related concepts or events tend to be recalled in succession^5,6^. In addition, transitions between distinct memories evoke characteristic neural responses^7^, similar to the neural dynamics observed when continuous external experiences are segmented and organized into discrete event representations^8,9^. However, these findings are primarily derived from studies involving the recall of experimentally induced experiences, such as reading word lists or watching movies^5,7^, where task demands control the flow of thoughts. What are the cognitive and neural mechanisms underlying the naturally occurring dynamics of memory and future thinking in real life?

Insights into the processes driving the naturalistic flow of memory and future thinking can be gained through the framework of spontaneous thought. Spontaneous thought refers to thoughts that arise and unfold freely, without being constrained by deliberate cognitive control or attention-capturing salient stimuli^10^. These thoughts mostly consist of personally relevant retrospective and prospective memories^4,11^, supported by semantic knowledge^3^, and often reflect the individual’s real-life goals and current concerns^12,13^. In addition, spontaneous thoughts share neural correlates with memory recall and future thinking^1,14–16^, particularly involving the default network^17^ and the frontoparietal control network^18^. The default network, including the hippocampus, is activated when thoughts are spontaneously generated and maintained^19,20^, such as during moments of self-reported mind-wandering^21,22^. The control network is also activated and functionally coupled with the default network during these instances^21,23,24^, and is thought to exert top-down control to guide the trajectory of thoughts^10,25^.

Despite this extensive research on spontaneous thought, the neurocognitive processes underlying the natural transitions and trajectory of spontaneous memory and future thinking remain underexplored. Common experimental paradigms, such as retrospective reports^26,27^ and experience sampling^21,22,26^, ask participants to report their thoughts after periods of rest or at intermittent intervals, limiting their ability to track the uninterrupted flow of ongoing thoughts. To address this limitation, recent studies have increasingly used the think-aloud paradigm, where participants verbalize their thoughts in real time during rest^13,28–33^, providing a more continuous and detailed report of naturalistic thoughts^30^. These studies have shown that thought trajectories are often clustered, with thoughts staying semantically related until transitioning to new topics, which creates boundaries between thoughts^13,28,29^. Moreover, the variability or stability of thought trajectories has been linked to distinct mental states^33^ and individual differences in personality and mental health^29,32^. However, the think-aloud paradigm has rarely been combined with neural recording techniques^30^, leaving important questions unanswered about how the brain generates and responds to these transitions and variability in thoughts.

Here, we used the think-aloud paradigm with functional magnetic resonance imaging (fMRI) to investigate the neural correlates of dynamic transitions between thoughts in the flow of spontaneous memory and future thinking. Focusing on the brain’s default and control networks, we aimed to address the following questions: 1) What are the major organizing principles guiding transitions from one thought to the next? 2) What are the neural signatures of these thought transitions? and 3) How do brain networks interact to generate variable or stable thought trajectories? We collected think-aloud responses during 10-minute resting fMRI scans and segmented them into discrete thought units, each containing a single topic and thought category (e.g., episodic memory, future thinking). By analyzing transition probabilities and semantic similarity between consecutive thoughts, we found that semantic associations primarily guided transitions to related thoughts, although shared neurocognitive processes (i.e., thought categories) also played a role. Strong thought boundaries, characterized by semantic disconnections, activated the default network and adjacent control network areas, resulting in distributed activation patterns similar to those observed at boundaries between external events^7^. Finally, interactions between the default and control network regions shaped the overall semantic structure of thought trajectories. Specifically, stronger functional connectivity within the default network subsystem including the hippocampus predicted greater semantic variability in thoughts, while stronger connectivity between the default and control networks was associated with reduced variability. Together, our findings highlight the central role of the default and control networks in organizing the natural transition dynamics and structure of the unconstrained stream of spontaneous memory and future thinking.

## RESULTS

### Content and distribution of thoughts

We first examined the content and distribution of various types of thoughts reported during the think-aloud fMRI session. Participants verbally described their stream of spontaneous thoughts for 10 minutes without interruption. Independent annotators manually segmented these responses into individual thought units based on changes in topic or category of thought (Fig. 1a). The identified categories were: current state including sensations and feelings (e.g., “I feel some breeze.”), semantic memory about the world or other people (e.g., “Baltimore’s pretty cool.”), semantic memory about oneself (e.g., “I’m a senior now.”), episodic memory (e.g., “I was walking around earlier with my boyfriend.”), imagining or planning the future (e.g., “I got to go to the grocery store.”), and other thoughts not fitting into the listed categories. Each thought unit was also assigned a topic label summarizing the content of the thought. Fig. 1d visualizes the most frequent topic labels for each thought category, aggregated across all participants.

**Fig. 1.**
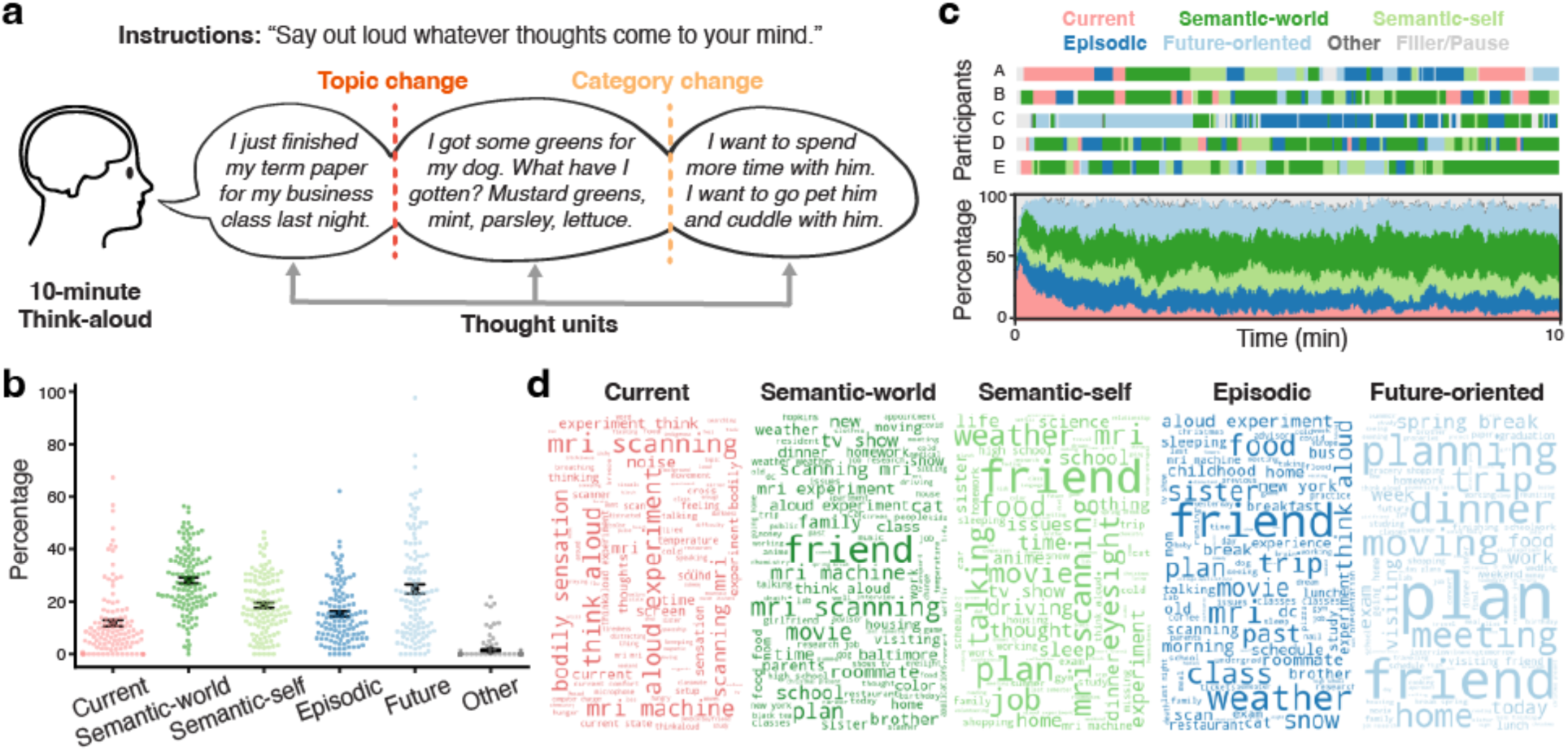
Think-aloud verbal responses. **a.** Participants verbally described their spontaneous flow of thoughts for 10 minutes inside the MRI scanner. Their speeches were transcribed and manually segmented into individual thought units, with each thought unit containing a single topic and corresponding to one of the following categories: current state, semantic memory about the world or other people, semantic memory about oneself, episodic memory, imagining or planning the future, and other uncategorized thoughts. **b.** Percentages of different thought categories among all thought units within each participant. Each colored dot represents an individual participant (N = 118 for all categories). Black circles indicate the mean across participants within each category. Error bars show the SEM across participants. **c.** Temporal distribution of different thought categories within the 10-minute think-aloud session. The upper panel shows the distribution of thought categories for five example participants. The lower panel shows the percentages of different thought categories averaged across participants for each time point (1 TR = 1.5-second window). **d.** Word clouds showing common topics for each major thought category. Topic labels were generated by the annotators who segmented the think-aloud responses. The 100 most frequent topic labels from the data combined across all participants are visualized using the WordCloud Python package (version 1.9.3). More frequent topics are shown in larger fonts.

Participants generated an average of 54.5 thoughts (SD = 19.9, range 19–118), producing an average of 1368.3 words (SD = 376.2, range 368–2268) excluding filler utterances (e.g., “Um, what else.”). Consistent with prior studies^4,34^, internally oriented thoughts involving memory and future thinking comprised the majority of spontaneous thoughts (M = 86.8%, SD = 13.6; Fig. 1b). Among these, semantic memory about the world/others was the most frequently reported (M = 28.1%, SD = 12.4), followed by future thinking (M = 24.8%, SD = 19.6), semantic memory about oneself (M = 18.6%, SD = 11.0), and episodic memory recall (M = 15.3%, SD = 11.2). On average, 11.8% of thoughts (SD = 13.3) described current states associated with performing the think-aloud task in the MRI scanner. Only 1.4% of thoughts (SD = 4.1) could not be categorized into one of the five major categories, confirming that our thought categorization scheme effectively captured the content of the think-aloud responses. For descriptive statistics related to each thought category, including mean duration, word count, speech rate, and streak length, see Supplementary Table 1.

The temporal distribution of thought categories over the 10-minute think-aloud session showed considerable individual variability (Fig. 1c, upper panel). To examine the group-level temporal distribution, we computed the proportion of participants who reported each thought category in each 1-TR (1.5 second) time window (Fig. 1c, lower panel). The thought categories were generally evenly distributed throughout the session, except that participants disproportionately reported thoughts describing the current state at the beginning of the scan. Specifically, current states comprised 57.6% of the first thoughts reported, suggesting that participants’ attention was initially captured by the salient external environment (i.e., being in the MRI scanner) before internally oriented thoughts emerged.

### Brain activation for different thought categories

We next examined brain activation associated with different categories of thoughts. First, we conducted a whole-brain analysis to identify the brain areas recruited during spontaneous memory recall and future thinking. For each cortical parcel from the Schaefer 400-parcel atlas^35^, we performed paired-samples *t*-tests comparing the mean activation of each of the four internally oriented thought categories against the current state category. The resulting group-level contrast maps are shown in Figs. 2a and 2b (Bonferroni corrected, *p* < .05). Consistent with prior findings^16,36^, the medial and lateral parietal cortices within the default network were more strongly activated during the description of internally oriented thoughts compared to the current state. This default network activation was more pronounced during episodic recall and future thinking (Fig. 2b) compared to describing generic semantic memory (Fig. 2a), highlighting its involvement in mental time travel and constructive simulation^17,37,38^. In contrast, the temporo-parietal junction, which overlaps with the salience/ventral attention network, was more strongly activated during the current state compared to the other categories. For the list of all suprathreshold parcels from each contrast, see Supplementary Tables 2-5.

**Fig. 2.**
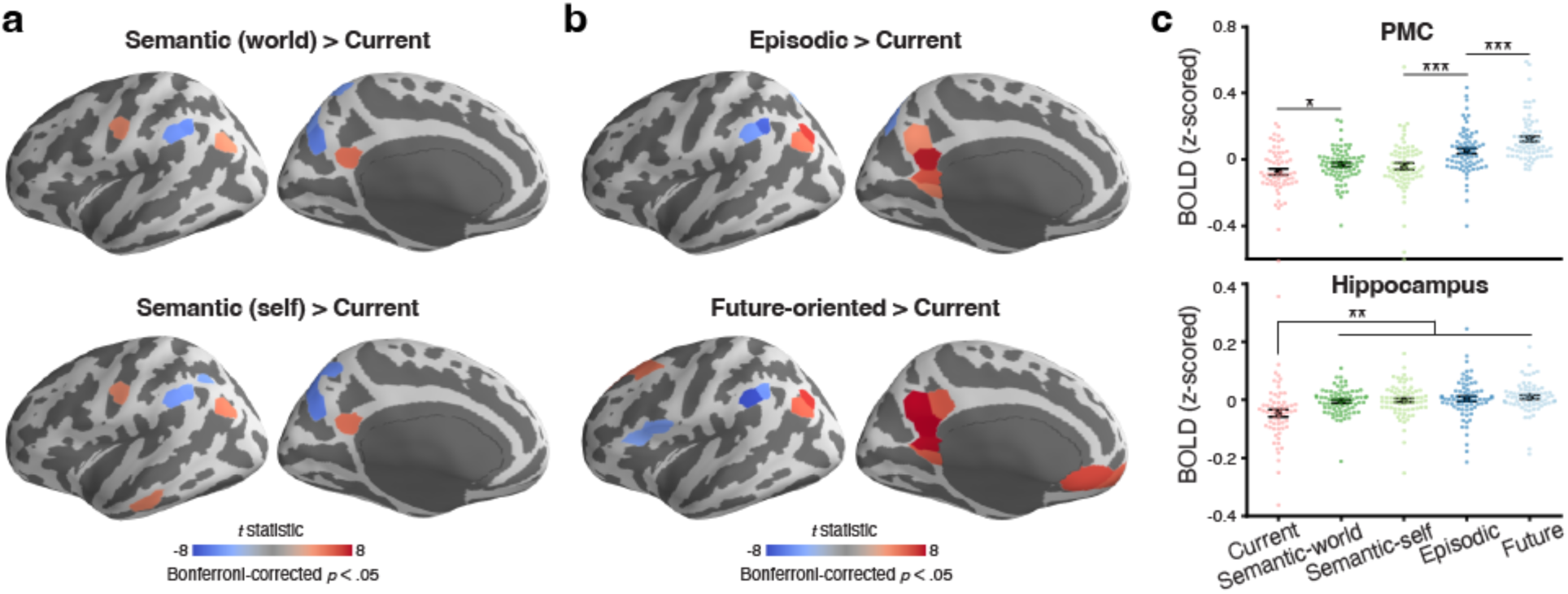
Univariate activation during memory recall and future thinking. **a.** Whole-brain *t*-statistic maps of cortical parcels showing higher or lower activation while describing semantic memory about the world or other people (top) or about oneself (bottom), compared to describing the current state. **b.** Whole-brain *t*-statistic maps of cortical parcels showing higher or lower activation while describing episodic memory (top) or future-oriented thoughts (bottom), compared to describing the current state. In both **a** and **b**, the *t*-statistic maps are displayed on the lateral (left) and medial (right) surfaces of the left hemisphere of the inflated fsaverage6 template brain. Parcels with significantly higher activation compared to the current state are shown in red, while those with significantly lower activation are shown in blue. The statistical significance of each contrast (*p* < .05) was Bonferroni corrected across the 400 parcels in the Schaefer atlas^35^. Supplementary Tables 2-5 provide the lists of suprathreshold parcels from both hemispheres. **c.** Mean blood oxygenation level-dependent (BOLD) signal for each thought category in the posterior medial cortex (PMC; top) and the hippocampus (bottom). Each colored dot represents an individual participant (N = 62, 75, 72, 73, and 72 for current, semantic-world, semantic-self, episodic, and future categories, respectively). Black circles indicate the mean across participants within each category. Error bars show the SEM across participants. \**p* < .05, \*\**p* < .01, \*\*\**p* < .001 (uncorrected).

Additionally, we examined the activation levels for different thought categories in two subregions of the default network: the posterior medial cortex (PMC) and the hippocampus (Fig. 2c). Both regions have been frequently implicated in memory retrieval, future thinking, and the generation of spontaneous thoughts^16,36^. Mean activation significantly varied across different thought categories in both PMC (*F*(4,224) = 22.72, *p* < .0001, *η*_p_^2^ = .29) and the hippocampus (*F*(4,224) = 5.28, Greenhouse-Geisser corrected *p* = .0017, *η*_p_^2^ = .09). In PMC, all internally oriented thought categories, except for semantic memory about oneself, showed higher activation compared to the current state (*t*s > 2.02, *p*s < .05, Cohen’s *d*s > .37). Mirroring the whole-brain analysis results, episodic recall and future thinking showed higher activation than semantic memory about oneself or the world (*t*s > 4.07, *p*s < .0002, Cohen’s *d*s > .65). Among these, future thinking activated PMC the most, with activation greater than episodic recall (*t*(70) = 3.73, *p* = .0004, Cohen’s *d* = .61, 95% CI = [.04, .12]). In the hippocampus, all internally oriented thought categories showed higher activation compared to the current state (*t*s > 2.7, *p*s < .01, Cohen’s *d*s > .50). However, there were no significant differences between the internally oriented thought categories themselves (*t*s < 1.91, *p*s > .06, Cohen’s *d*s < .32).

### Transitions between thoughts

An important characteristic of the continuous flow of thoughts is that the mind continually moves from one thought to another^3,28,39^, switching between topics and categories (Fig. 1c). What principles underlie the dynamics of these thought transitions? Are there specific mental states that trigger spontaneous memory recall and future thinking? One possibility is that a thought may be evoked by another thought sharing similar neurocognitive processes, such as when memory retrieval is more likely to follow previous memory retrieval than the encoding of new information^40–42^. In this context, a thought is likely to be followed by another from the same category, leading to temporally clustered thought categories. To test this idea, we employed a Markov chain approach following prior studies^29,43,44^, and computed transition probabilities across the six thought categories including the “other” category (Fig. 3a). We calculated these probabilities between individual sentences rather than thought units to avoid bias that arises from using category transitions to define thought unit boundaries. Consistent with our prediction, the probability of a thought category transitioning to itself (i.e., the diagonal values of the transition probability matrix) was higher than expected by chance in all thought categories except for the “other” category (*t*s > 12.78, *p*s < .0001, Cohen’s *d*s > 1.23).

**Fig. 3.**
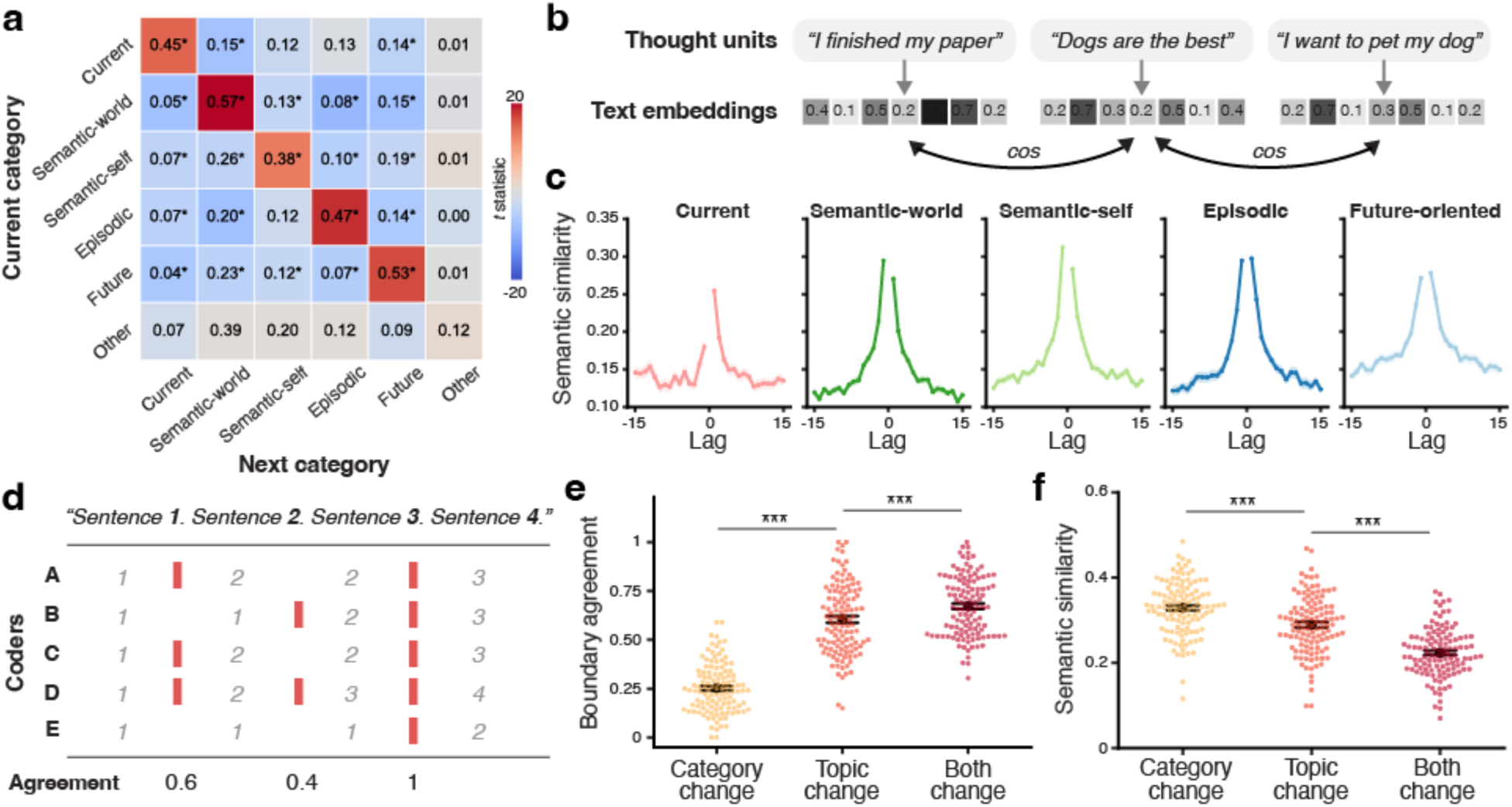
Transitions between thoughts. **a.** Sentence-level transition probability between different thought categories. The rows and columns of the matrix represent the current and next categories, respectively. The numbers in the matrix indicate transition probabilities for each category pair, averaged across participants. The colormap of the matrix indicates the *t*-statistics from one-sample *t*-tests against the chance probability (i.e., the overall proportion of the next category among all sentences within each participant). Transitions that occur more frequently than chance are shown in red, while those that occur less frequently than chance are shown in blue. \**p* < .05 (Bonferroni corrected). **b.** Measuring semantic similarity between thought units. Each thought unit was converted to a text embedding vector using the Sentence Transformers Python module (version 2.2.0). Semantic similarity between thoughts was defined as the cosine similarity between their embedding vectors. **c.** Semantic similarity as a function of the temporal distance from a target thought unit in each thought category. Lags are measured in units of thought, with lag = 0 representing the target thought. Negative and positive lags indicate thoughts that occurred before and after the target thought, respectively. Solid lines indicate the mean across participants (N = 98, 117, 113, 113, and 112 for current, semantic-world, semantic-self, episodic, and future categories, respectively). Shaded areas indicate the SEM across participants. **d.** Measuring thought boundary agreement scores from think-aloud transcripts. Independent coders assigned the same numbers to consecutive sentences/clauses describing a single thought. Thought boundaries (red bars) were detected when the thought identification numbers changed. Boundary agreement scores were defined as the proportion of coders who identified each moment as a thought boundary. **e.** Mean boundary agreement scores for different types of thought transitions. **f.** Mean semantic similarity between pre- and post-boundary thoughts for different types of thought transitions. In both **e** and **f**, each colored dot represents an individual participant (N = 117, 118, and 117 for category change, topic change, and both change conditions, respectively). Black circles indicate the mean across participants within each transition type. Error bars show the SEM across participants. \*\*\**p* < .001 (uncorrected).

Another potential major organizing factor in the chain of thoughts is semantic relations. Models of episodic and semantic memory search^5,45,46^ and spontaneous thought^13,39^ suggest that shared meanings can cue semantically associated thoughts. To test this, we measured semantic similarity between thought units using a natural language processing technique^28,31,33^, defining it as the cosine similarity between text embedding vectors representing each thought (Fig. 3b). Supporting the semantic association hypothesis, we found that a thought was semantically more similar to its immediate consecutive thoughts (lags −1 and 1) than to more temporally distant thoughts (lags −15 and 15) across all thought categories (*t*s > 9.98, *p*s < .0001, Cohen’s *d*s > 1.22; Fig. 3c). The semantic association between consecutive thoughts was particularly stronger for internally oriented thought categories including memory and future thinking, compared to the current state category (*t*s > 4.41, *p*s < .0001, Cohen’s *d*s > .57).

If both shared neurocognitive states (i.e., thought categories) and semantic associations affect transitions between thoughts, which factor has a greater impact? To address this question, we compared thought boundaries involving category changes with those involving topic changes (Fig. 1a) in terms of their perceived disconnectedness. If semantic associations play a more significant role in the flow of thoughts, then changes in topics (e.g., shifting from episodic recall about a term paper to episodic recall about a dog) will be perceived as stronger boundaries than changes in general thought categories (e.g., shifting from episodic recall about a dog to semantic memory about the dog), and vice versa. To independently measure the perceived strength of boundaries between thoughts, we had a separate group of human coders read the think-aloud transcripts and identify moments when one thought transitioned to another (Fig. 3d). Critically, they were instructed to use their best subjective judgment based on any criteria and were not specifically told to consider changes in thought categories or topics. The measure of boundary strength was boundary agreement scores, computed as the proportion of coders who identified each moment as a thought boundary.

The results suggested that semantic associations may play a more crucial role than thought categories in defining thought boundaries. Boundary agreement scores varied across different types of thought boundaries (Fig. 3e; *F*(2,230) = 501.48, *p* < .0001, *η*_p_^2^ = .81), with higher agreement observed at boundaries involving both topic and category changes (*t*(115) = 28.67, *p* < .0001, Cohen’s *d* = 3.00, 95% CI = [.39, .45]), or topic changes alone (*t*(116) = 24.06, *p* < .0001, Cohen’s *d* = 2.27, 95% CI = [.32, .38]), compared to those involving only category changes. This pattern was mirrored in the semantic similarity between pre- and post-boundary thoughts: semantic similarity was lower at boundaries involving both topic and category changes (*t*(115) = 15.29, *p* < .0001, Cohen’s *d* = 1.74, 95% CI = [.09, .12]) and topic changes alone (*t*(116) = 5.57, *p* < .0001, Cohen’s *d* = .61, 95% CI = [.03, .05]), compared to those involving only category changes (Fig. 3f). Furthermore, boundary agreement was negatively correlated with semantic similarity between consecutive thoughts within each participant (mean *r* = -.35, SD = .16; one-sample *t*-test against zero: *t*(116) = −23.75, *p* < .0001, Cohen’s *d* = 2.20, 95% CI = [-.37, -.32]), confirming that changes in semantic content critically influenced thought boundary perception.

### Neural responses at major thought transitions

Although internally oriented thoughts generally transition to semantically related ones, shifts to unrelated topics occasionally occur, creating salient boundaries^28,47^. What are the neural signatures of these prominent boundaries between thoughts? While neural responses at event boundaries driven by changes in external stimuli have been studied extensively^48–50^, internally-driven boundaries between mental contexts have rarely been investigated^7,51^. To characterize the neural responses at boundaries between thoughts, we focused our analysis on the strongest boundaries, defined by a boundary agreement score of 1 (pre- and post-boundary thought semantic similarity: M = .19, SD = .08). Among these, 80.7% involved transitions to one of the four memory/future thinking categories. Supplementary Table 6 provides a breakdown of the percentages for specific thought category pairs that preceded and followed the strong thought boundaries.

We began by identifying the brain areas activated at strong thought boundaries. We performed a whole-brain univariate analysis, contrasting the average activation during boundary periods with that during non-boundary periods (Fig. 4a). A boundary period was defined as a 6-second window following the offset of a pre-boundary thought. A non-boundary period was defined as a 6-second window in the middle of a thought lasting longer than 15 seconds.

**Fig. 4.**
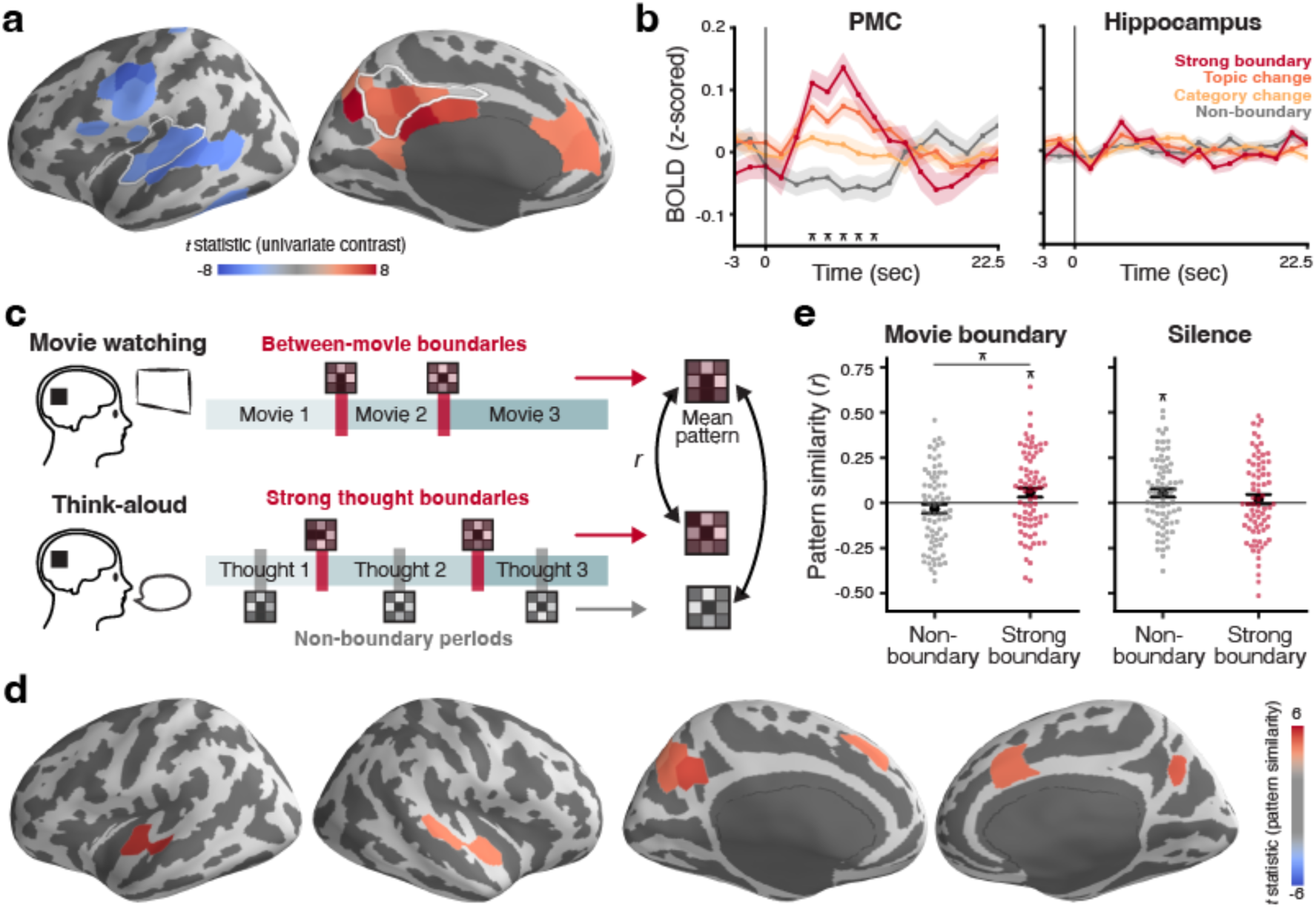
Neural responses at thought boundaries. **a.** Whole-brain *t*-statistic map of the univariate contrast between strong boundary (boundary agreement = 1) and non-boundary periods. Parcels with significantly higher activation during strong boundary periods compared to non-boundary periods are shown in red, while those with significantly lower activation are shown in blue. Statistical significance (*p* < .05) was Bonferroni corrected across all parcels. White outlines indicate the auditory cortex and the PMC ROIs, respectively. **b.** Mean PMC (left) and hippocampus (right) activation time courses aligned at different types of thought boundaries. Time zero for the non-boundary condition represents the middle of thoughts longer than 15 seconds. For other conditions, time zero represents the offset of the pre-boundary thought. Solid lines indicate the mean across participants (N = 75 for all conditions). Shaded areas indicate the SEM across participants. Asterisks above the x-axis indicate time points where activation for strong boundaries is significantly higher than non-boundaries after Bonferroni correction (*p* < .05). **c.** Boundary pattern similarity analysis. For each region, we computed the mean activation pattern of between-movie boundaries from the movie-watching phase of our prior study^7^. This template pattern was correlated with the mean activation patterns of strong thought boundaries (red bars) and non-boundary periods (gray bars) during think-aloud. **d.** Whole-brain *t*-statistic map of boundary-specific pattern similarity. Parcels are shown in red if their between-movie boundary patterns were more similar to their strong thought boundary patterns than to the non-boundary patterns. The map is masked to only include areas that showed positive correlations between the between-movie boundary patterns and the strong thought boundary patterns. Statistical significance (*p* < .05) was Bonferroni corrected across all parcels. **e.** Boundary pattern similarity in PMC. The think-aloud strong thought boundary and non-boundary patterns were correlated with the mean activation patterns of between-movie boundary periods (left panel) or silent periods (right panel) from the movie watching phase^7^. Each colored dot represents an individual participant (N = 75 for all conditions). Black circles indicate the mean across participants. Error bars show the SEM across participants. \**p* < .05 (uncorrected).

Greater activation during boundary periods, compared to non-boundary periods, was observed primarily in the medial frontal and parietal areas of the default network and control network. In contrast, greater activation during non-boundary periods was observed in the lateral frontal areas associated with speech generation and areas around the auditory cortex, reflecting the effect of a temporary pause in speech at thought boundaries. For the list of all suprathreshold parcels, see Supplementary Table 7.

We further examined the activation time course in the PMC and hippocampus ROIs for different types of thought boundaries (Fig. 4b). PMC showed significant activation from 4.5 to 10.5 seconds following strong thought boundaries, compared to the non-boundary time course aligned to the middle of thoughts (*t*s > 4.27, *p*s < .0001, Cohen’s *d*s > .66). The boundary responses in PMC were also scaled with the strength of thought boundaries. Boundaries involving only topic changes (boundary agreement M = .60, SD = .18) evoked weaker responses compared to the strong boundaries with agreement scores of 1. Boundaries involving only thought category changes (boundary agreement M = .25, SD = .13) resulted in even weaker responses. The hippocampus showed a slightly higher response at 4.5 seconds following strong boundaries compared to non-boundaries, which did not reach statistical significance (*t*(74) = 1.96, *p* = .054, Cohen’s *d* = .37, 95% CI = [-.00, .08]).

Next, we analyzed the distributed activation patterns at strong thought boundaries. In a prior study^7^, we identified a distinctive activation pattern associated with major mental context transitions within the default network and the adjacent control network, particularly around PMC. Specifically, we observed highly similar activation patterns at boundaries between different movies while participants watched a series of films. These consistent patterns also reappeared at boundaries between memories of the movies during continuous verbal recall. We predicted that this major mental context transition pattern would generalize to strong thought boundaries during think-aloud sessions.

To test this, we conducted a whole-brain pattern similarity analysis (Fig. 4c). For each cortical parcel, we correlated the mean activation pattern at strong thought boundaries during think-aloud sessions with the mean activation pattern at between-movie boundaries from the movie watching phase of our prior study^7^. We also correlated the mean non-boundary activation pattern during think-aloud with the same between-movie boundary pattern. As predicted, the major mental context transition pattern was observed in parcels within and around PMC. Figure 4d illustrates these parcels, where 1) strong thought boundary patterns were positively correlated with between-movie boundary patterns, and 2) this correlation was greater than the correlation between non-boundary patterns and between-movie boundary patterns.

Supplementary Figure 1 shows separate whole-brain maps of positive pattern similarities between thought boundaries and movie boundaries (Suppl. Fig. 1a) and significant differences between strong thought boundaries and non-boundaries (Suppl. Fig. 1b). Similar results were observed within the PMC ROI (Fig. 4e, left panel), showing a positive correlation between the strong thought boundary pattern and the between-movie boundary pattern (*t*(74) = 2.22, *p* = .029, Cohen’s *d* = .26, 95% CI = [.01, .11]). This correlation was also greater than the correlation between the non-boundary pattern and the between-movie boundary pattern (*t*(74) = 2.29, *p* = .025, Cohen’s *d* = .41, 95% CI = [.01, .17]).

Is this thought transition pattern simply driven by pauses in speech at boundaries? Strong thought boundaries in the current study and between-movie boundaries in ref.^7^ share low-level auditory features, as both involve brief periods of silence. Indeed, parcels around the auditory cortex also showed a positive correlation between strong thought boundary patterns and between-movie boundary patterns (Fig. 4d). To rule out this possibility, we compared the activation patterns at strong thought boundaries with those during periods of silence in the auditory cortex and PMC ROIs. The silence pattern was derived from the movie-watching phase of ref.^7^ by averaging silent moments within the movie stimuli. In the auditory cortex (Suppl. Fig. 2), the silence pattern was positively correlated with the strong thought boundary pattern (*t*(74) = 4.85, *p* < .0001, Cohen’s *d* = .56, 95% CI = [.07, .18]) but negatively correlated with the non-boundary pattern (*t*(74) = −3.17, *p* = .002, Cohen’s *d* = .37, 95% CI = [-.10, -.02]), confirming that its thought boundary pattern was driven by the absence of sound. In contrast, in PMC (Fig. 4e, right panel), the silence pattern was not correlated with the strong thought boundary pattern (*t*(74) = .75, *p* = .46, Cohen’s *d* = .09, 95% CI = [-.03, .07]), but was positively correlated with the non-boundary pattern (*t*(74) = 2.39, *p* = .019, Cohen’s *d* = .28, 95% CI = [.01, .10]). Thus, the internally-driven boundary pattern in PMC is unlikely to be driven by pauses in speech.

### Thought structure and brain connectivity

So far, we have focused on transitions between immediately neighboring thoughts. However, the dynamics of thought can also be reflected in the overall semantic structure, including the relationships between temporally distant thoughts, such as the recurrence of similar topics over time. Indeed, individuals’ thought streams vary in how divergent or focused their content is^10,33^. What are the neural underpinnings of this variability or stability in thoughts? A prominent perspective on spontaneous thought hypothesizes that various large-scale brain networks play distinct roles in shaping the structure of internally oriented thoughts^10^. The medial temporal lobe subsystem of the default network may be responsible for generating variable thoughts, while the core default network subsystem likely constrains thoughts toward personally significant information^52^. The frontoparietal control network (FPCN) may interact with other networks to help sustain goal-relevant thoughts, thereby increasing thought stability^33^.

To test this idea, we explored the relationship between functional connectivity within and between large-scale brain networks and the overall semantic structure of think-aloud responses. The semantic structure was quantified using the average clustering coefficient of the semantic network of thoughts, where nodes represented individual thought units and edges represented the semantic similarity between these thoughts (Fig. 5a). Higher clustering coefficients indicated more stable and focused thought structures, while lower clustering coefficients indicated more variable and divergent thought structures (Fig. 5b). Figure 5c shows the distribution of average clustering coefficients across all participants (M = .18, SD = .05). For the functional connectivity analysis, we focused on Control Network B, Default Network A, and Default Network C as defined in the 17-network version of the Schaefer atlas^35^, with the hippocampus included in Default Network C (Fig. 5d). These networks correspond, respectively, to the FPCN, the core default network subsystem, and the medial temporal lobe subsystem of the default network, as outlined in ref.^10^.

**Fig. 5.**
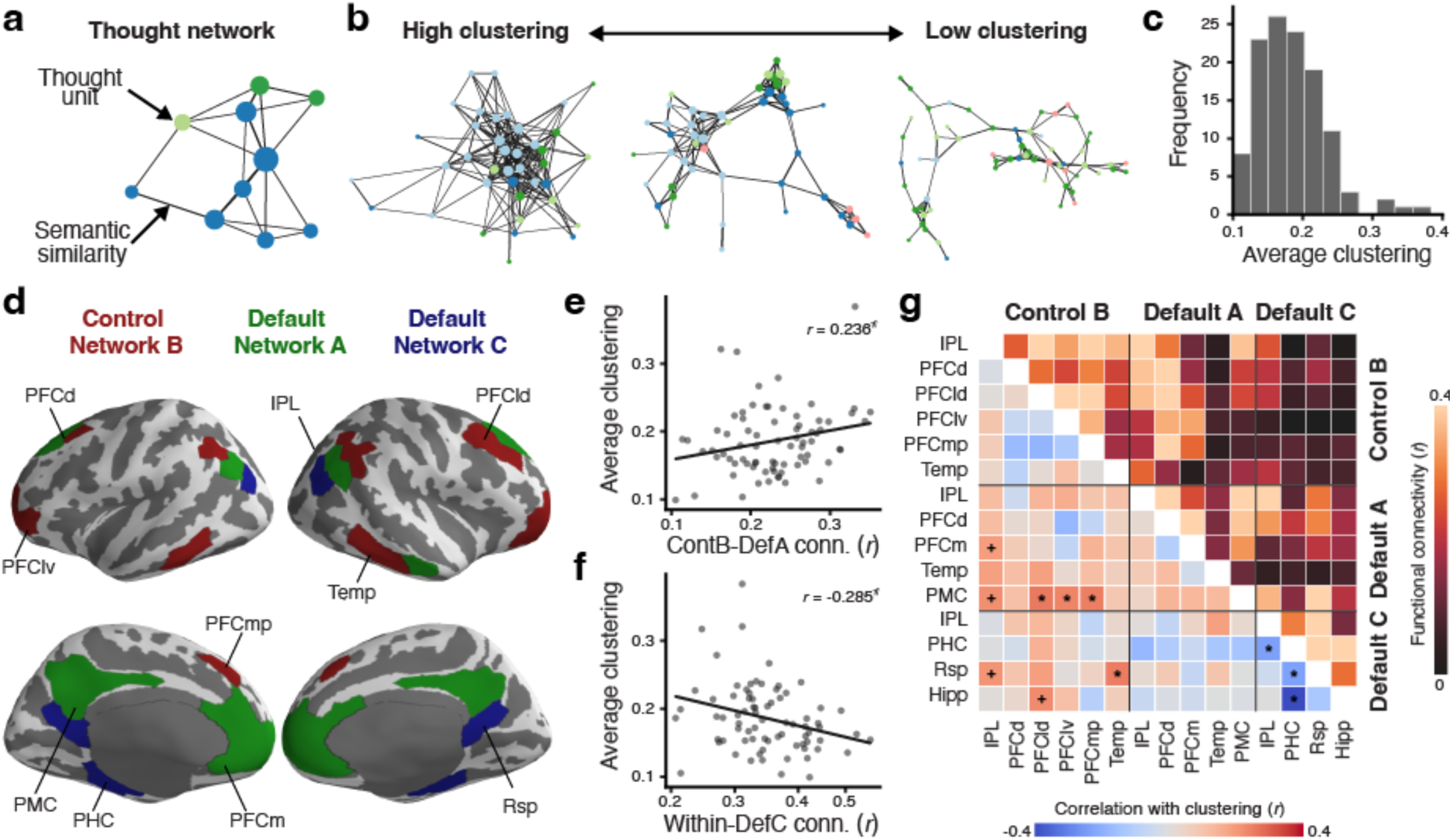
Thought network structure and functional connectivity. **a.** A thought network where nodes represent thought units, and edge weights represent semantic similarity between thoughts. **b.** Example thought networks with three different levels of clustering (average clustering coefficients = 0.37, 0.27, and 0.19). For visualization, edge weights were thresholded at a cosine similarity of 0.3. In both **a** and **b**, node size and edge thickness are proportional to normalized degree and edge weights, respectively. Different node colors represent different thought categories (pink = current; green = semantic-world; light green = semantic-self; blue = episodic; light blue = future-oriented). **c.** Distribution of the average clustering coefficients of thought networks generated from the think-aloud responses of all 118 participants included in the behavioral analyses. Thought network edge weights were thresholded at zero. **d.** Subregions of Control Network B (red), Default Network A (green), and Default Network C (blue) as defined in the 17-network version of the 400-parcel Schaefer atlas^35^. The subregions are displayed on the lateral (top) and medial (bottom) surfaces of the inflated fsaverage6 template brain. **e.** Pearson correlation between the average clustering coefficients of thought networks and the between-network functional connectivity between Control Network B and Default Network A. **f.** Pearson correlation between the average clustering coefficients of thought networks and the within-network functional connectivity in Default Network C. In both **e** and **f**, gray dots represent each of the 75 participants included in fMRI analyses. Solid lines represent the best-fitting regression lines. **g.** Mean functional connectivity between the subregions of the three brain networks of interest (upper triangle) and the correlation between the functional connectivity and thought network clustering coefficients (lower triangle). Hipp = hippocampus; IPL = interior parietal lobule; PFCd = dorsal prefrontal cortex; PFCld = lateral dorsal prefrontal cortex; PFClv = lateral ventral prefrontal cortex; PFCm = medial prefrontal cortex; PFCmp = medial posterior prefrontal cortex; PHC = parahippocampal cortex; PMC = posterior medial cortex; Rsp = retrosplenial cortex; Temp = temporal lobe. +*p* < .1, \**p* < .05 (uncorrected).

As expected, interaction between the control and default networks was associated with semantic stability in think-aloud responses. Specifically, functional connectivity between Control Network B and Default Network A was positively correlated with thought network clustering coefficients across participants (*r*(75) = .24, *p* = . 041, 95% CI = [.01, .44]; Fig. 5e). Functional connectivity between Control Network B and Default Network C was also numerically positively correlated with clustering coefficients, although this relationship did not reach statistical significance (*r*(75) = .13, *p* = . 256, 95% CI = [-.10, .35]). In contrast, within-network functional connectivity computed across the subregions of Default Network C was negatively correlated with thought network clustering coefficients (*r*(75) = -.29, *p* = . 013, 95% CI = [-.48, -.06]; Fig. 5f), supporting its role in generating thought variability^10^. There was no significant correlation between clustering coefficients and the connectivity between Default Network A and Default Network C (*r*(75) = -.04, *p* = . 729, 95% CI = [-.27, .19]).

Additionally, we performed a post-hoc exploratory analysis to identify specific pairs of subregions whose functional connectivity correlates with the overall semantic structure of thoughts (Fig. 5g, lower triangle). We found that connectivity between PMC in Default Network A and the lateral dorsal prefrontal, lateral ventral prefrontal, and medial posterior prefrontal cortex subregions in Control Network B was positively correlated with thought network clustering coefficients (*r*(75)s > .23, *p*s < .041). Connectivity between the temporal lobe subregion of Control Network B and the retrosplenial cortex in Default Network C was also positively correlated with clustering coefficients (*r*(75) = .25, *p* = .029, 95% CI = [.03, .45]). In contrast, within Default Network C, connectivity between the parahippocampal cortex (PHC) and the hippocampus, retrosplenial cortex, and inferior parietal lobule subregions was negatively correlated with thought network clustering coefficients (*r*(75)s < -.23, *p*s < .044). The strongest of these correlations was between PHC-hippocampus connectivity and clustering coefficients (*r*(75) = -.42, *p* = .0002, 95% CI = [-.59, -.21]), which survived Bonferroni’s correction for multiple comparisons.

## DISCUSSION

The current study investigated the neural mechanisms underlying the dynamic flow of spontaneous memory recall and future thinking. Using a think-aloud paradigm, where participants continuously verbalized their thoughts during resting fMRI scans, we captured neural responses specifically linked to natural transitions between thoughts and the semantic structure of thought trajectories. Within the flow of thought, primarily consisting of retrospective and prospective memories, transitions predominantly occurred between semantically associated thoughts. Notably, significant shifts in the semantic content of thoughts created boundaries between them, activating core posterior-medial areas of the default and control networks. These boundary responses generated distributed activation patterns comparable to those evoked by boundaries between external events. Furthermore, functional connectivity within and between the default and control networks predicted the overall semantic variability and stability of thought trajectories, highlighting the crucial role of these large-scale networks in shaping the dynamics of spontaneous memory and future thinking.

The think-aloud paradigm enabled us to identify brain regions involved in naturally arising memory and future thinking. Compared to thoughts focused on current feelings and sensations, these internally oriented thoughts—particularly episodic recall and future imagination—activated the default network including the hippocampus, medial frontal cortex, lateral parietal cortex, and PMC. This activation did not systematically correlate with behavioral measures such as thought duration, word count, or speech rate (Supplementary Table 1), suggesting that it reflects deeper cognitive processes rather than superficial features of speech or thought production. Prior studies using more controlled tasks have also implicated a similar set of regions in episodic memory retrieval^15,16^, mental simulation^1^, and self-referential thinking^19,43^, further supporting their role in constructing internal narratives^53^. In addition to the default network, spontaneous thought generation is known to engage broader neural systems, including regions involved in cognitive control^21,23,24^. However, we did not observe notable activation in the control network when compared to the current state category. While the current state category primarily captured thoughts about the immediate environment, the process of consciously accessing and verbalizing them within a continuous stream may demand a similar level of cognitive control as memory and future thinking. This overlap could have diminished the contrast between current state and internally oriented thought categories.

We found that both shared neurocognitive processes and semantic connections guide the transitions of spontaneous thoughts. Specifically, 1) thoughts tend to transition within the same thought category, and 2) consecutive thoughts show higher semantic similarity than temporally distant ones. This reflects a tendency for thoughts to remain stable for a period before switching to a new one, consistent with prior research describing the locally clustered structure of thought trajectories^3,28,54^. Additionally, topic transitions elicited stronger boundary perceptions and greater cortical activation than thought category transitions, suggesting that semantic connections play a more dominant role in driving spontaneous thought transitions. This finding reinforces the longstanding view that semantic associations provide an organizing framework for internal representations and can serve as retrieval cues^3,5,39,46^. That said, it is worth noting that thought category and semantic content may not be entirely separable. For example, in our study, the current state category predominantly involved semantic content related to the MRI scanning environment (Fig. 1d). A previous study^27^ has also reported correlations between temporal dimensions (e.g., past, future) and content-related dimensions (e.g., people, images) in spontaneous thought. Future research could further investigate how different thought categories and semantic content interact to influence transitions between thoughts.

At strong thought boundaries marked by prominent shifts in thought content, midline default and control network regions are recruited, generating distributed activation patterns similar to those observed at externally-driven event boundaries. Neural responses to stimulus-driven boundaries between external events have been extensively studied in the fields of perception and memory, as they reveal how the brain segments and encodes continuous experiences into discrete events^8,49,50^. However, responses to boundaries created by internal mental context transitions remain largely unexplored^51,55^. In a rare prior study^7^, we demonstrated that boundaries between memories of different movies during continuous narrated recall elicit stereotyped activation patterns in PMC and nearby areas, similar to those seen at stimulus-driven movie boundaries during the initial viewing. The current study replicates and expands on these findings, applying them to boundaries between spontaneous internal narratives, which encompass broader semantic topics and exhibit more unconstrained dynamics. Our findings suggest that the boundary responses in the posterior medial areas represent a generalized signal of mental context transitions. This signal likely reflects internal task-switching demands^56,57^, which arise at the end of a thought to resolve competition among upcoming thoughts, allowing one to dominate conscious attention. Supporting this idea, the regions with heightened activation at thought boundaries overlapped with posterior-medial areas of the control network (Fig. 4a, Supplementary Table 7). These areas are known to play a key role in top-down cognitive control during task set changes^58,59^. However, this interpretation relies on reverse inference^60^, and further research is needed to fully understand the nature of cortical boundary responses in spontaneous thought.

Despite robust cortical responses, we did not observe significant hippocampal activation at major thought boundaries. This was unexpected given the hippocampus’s well-established role in spontaneous thought generation^10,47,61^ and mental time travel^36^. Moreover, the hippocampus is consistently activated at externally-driven boundaries between naturalistic events, supporting the successful encoding of these events^49,62,63^. Why, then, does the hippocampus not respond to thought boundaries? One possible explanation is the continuous demand for memory retrieval and thought generation inherent in the think-aloud task. This may lead to sustained hippocampal activation throughout most of the session, masking any responses to thought boundaries, if they exist. Indeed, the hippocampus showed consistently higher activation for memory and future thinking categories compared to the current state category (Fig. 2c). In contrast, during tasks that primarily involve encoding new external events, such as watching movies, the hippocampus may respond specifically to event boundaries by transiently retrieving the just-concluded event^64^. Thus, hippocampal responses differ between externally and internally driven mental context boundaries, despite similar cortical activation patterns.

Our findings demonstrate the role of functional connectivity within and between large-scale brain networks in shaping the semantic structure of spontaneous thought trajectories. This connectivity provides a potential neural basis for individual differences in thought dynamics, ranging from fleeting and freely flowing to more controlled and sustained patterns^65^. Consistent with the dynamic framework of spontaneous thoughts^10^, stronger interactions within the medial temporal lobe subsystem of the default network (DN_MTL_) were linked to greater thought variability, positioning it as a source of variability through associative cueing and pattern completion^39^. In contrast, increased coupling between the control network and the core subsystem of the default network (DN_core_) was associated with thought stability, confirming the role of the control network in constraining thoughts toward goal-relevant content^52^. These findings also align with creativity research, which suggests that the default network facilitates divergent idea generation, while the control network monitors and evaluates these ideas for goal-relevance^66^. However, connectivity between DN_MTL_ and DN_core_ did not correlate with thought structure, despite the DN_core_’s proposed role in automatically constraining thoughts toward salient internal information^10^. This may be because automatic constraints can either increase or decrease thought variability depending on its nature^52^. For example, automatic constraints may reduce variability during rumination, when individuals fixate on negative thoughts or emotions. Conversely, they may increase variability by triggering shifts to salient but irrelevant thoughts when attempting to focus on goal-relevant topics.

Although the think-aloud paradigm has significantly advanced our ability to capture the neural dynamics of the continuous flow of thoughts, investigating the fully unconstrained and spontaneous nature of real-world thought remains challenging. Spontaneous thoughts are deeply intertwined with real-life contexts and actions^67^, and the fixed setting of verbalizing thoughts in an MRI machine may restrict their natural contents and flow. Moreover, the presence of experimenters and the awareness of being recorded can lead to self-censorship or over-explanation, as indicated by the higher percentage of general semantic descriptions in our data (Fig. 1b, Semantic-world) compared to prior reports^11^. Even without social influences, the very act of consciously accessing and verbalizing thoughts could potentially alter the trajectory of spontaneous thinking^68^. Future research may explore how metacognition^69^ and the generation of external or internal speech^70^ affect the structure and transition dynamics of spontaneous memory and future thinking.

In conclusion, our study uncovers the cognitive and neural processes underlying the spontaneous flow of retrospective and prospective memory, bridging the fields of memory and spontaneous thought. Specifically, the default and control networks play a crucial role in thought transitions, and their interactions shape the overall structure of thought trajectories.

Understanding these dynamics aids in decoding resting state neural activity^26,27^, which has been widely used to explore the neural underpinnings of both clinical conditions and basic cognitive processes. Furthermore, the unfolding of spontaneous thoughts over time reflects the organization of naturalistic thought and predicts individual differences in personality^32,43,54^, mental health^20,29,71^, and well-being^72,73^. By investigating the brain mechanisms driving thought dynamics, our findings offer insights for future research aimed at developing novel biomarkers and interventions for psychiatric disorders, as well as promoting creativity and emotional well-being.

## METHODS

The current study adheres to ethical regulations governing research involving human participants. All experimental procedures were in accordance with protocols approved by the Institutional Review Boards of Johns Hopkins Medicine and Homewood.

### Participants

We recruited 126 healthy participants from the Johns Hopkins University community (76 females, age 18 – 40 years, mean age 23.7 years). All participants were right-handed native English speakers and reported normal hearing as well as normal or corrected-to-normal vision. Informed consent was obtained following procedures approved by the Johns Hopkins Medicine Institutional Review Board. Participants received monetary compensation for their time.

Of the 126 participants initially recruited, 8 were excluded from both behavioral and fMRI data analyses for the following reasons: poor quality of speech audio recordings (5 participants), scanning interruptions due to technical issues (2 participants), and failure to adhere to instructions (1 participant). An additional 43 participants were excluded from the fMRI data analysis due to: excessive head motion, defined as a mean framewise displacement greater than .5 mm (39 participants); anomalies in brain structure (2 participants); technical issues related to visual presentation using the projector (1 participant); and an unidentified artifact in the MRI data (1 participant). Consequently, 118 participants were included in behavioral data analyses (73 females, age 18 – 39 years, mean age 23.4 years), and 75 participants were included in fMRI data analyses (43 females, age 18 – 36 years, mean age 23.4 years).

### Study procedures

Participants completed a single 10-minute think-aloud session in the MRI scanner, during which they verbally described their spontaneous flow of thoughts (Fig. 1a). They were instructed to continuously speak out loud whatever thoughts came to their minds, including but not limited to memories of past events, plans for the future, or any bodily sensations, sights, sounds, or other feelings that captured their attention during the experiment. Participants were instructed to let their thoughts flow freely and not force themselves to stick to a single topic. They were asked not to entertain the experimenter or explain their thoughts by providing background information. Participants were allowed to refrain from verbalizing private thoughts if they did not wish them to be heard. Instead, they were instructed to briefly mention the topic of the thought and state that they did not want to share it (e.g., “It reminded me of my parents but I will not talk about it”).

Participants began speaking when the word “begin” appeared in white text on a gray screen. After 2 seconds, the “begin” message disappeared, and a white fixation cross was presented at the center, remaining on the screen throughout the task. Participants were instructed to keep their eyes open and look at the fixation cross. However, they were not required to maintain fixation on the cross for the entire task, and their eye movements were not monitored. The visual stimuli were presented on the screen located behind the magnet bore and viewed via an angled mirror. Participants’ speech was recorded using an MR-compatible microphone (FOMRI II; Optoacoustics Ltd.).

In all but two participants, various tasks unrelated to the current study were performed following the think-aloud task. The remaining two participants performed the think-aloud task at the end of the scanning session, following the unrelated tasks. The unrelated tasks included listening to audio stories, generating word chains, watching screen recording videos, browsing the web, and verbally recalling memories. Different combinations of these tasks were performed in each scanning session, and the results from these tasks will be reported elsewhere.

After the fMRI scanning session, participants received a link to a battery of online questionnaires asking about their personality traits, mental health, and demographic information. They were instructed to complete the questionnaires within two days following the fMRI session. Sixty-nine out of the 126 participants completed the questionnaires. Results from the questionnaires will be reported elsewhere.

### Behavioral data preprocessing

The audio recording of each participant’s think-aloud response was transcribed either manually or automatically using Whisper (Large-v2 model; OpenAI) and subsequently manually corrected. Each transcript was segmented into sentences, and timestamps were identified for the beginning and end of each sentence. Transcribed sentences that ended before the beginning of the scan or began after the end of the scan were excluded from analysis.

The transcripts were further processed by 15 independent human annotators. Each transcript was handled by a single annotator, with each annotator processing an average of 7.9 transcripts (range: 1 – 35). The annotators manually categorized each transcribed sentence into one of the following seven categories: 1) current state, action, or sensation during the experiment, 2) general knowledge or opinion about the world or other people (semantic-world), 3) general knowledge or opinion about oneself (semantic-self), 4) memories of past events in specific times and places (episodic), 5) imagining or planning the future (future-oriented), 6) filler utterances without specific content (e.g., “Um, what else.”), and 7) other utterances that cannot be categorized as any of the above categories.

The annotators also identified the topic of the thought described in each sentence and provided a short label of the topic (e.g., MRI scanning, cold weather, flu shot). To ensure consistency in topic labeling, the annotators were instructed to use the same label if the same topic was repeated within a transcript. In case either the thought category or the topic of the thought changed within a single sentence, the sentence was further broken down into multiple clauses, ensuring that no segment was coded as having more than one category or topic. Consecutive sentences or clauses with the same category describing the same topic were combined to form a single ‘thought unit’ (Fig. 1a).

### Thought category transition probability

Within each participant, we calculated transition probabilities between thought categories. These probabilities were computed between individual sentences rather than coarser thought units to avoid bias. Consecutive thought units were biased against belonging to the same category because transitions between thought categories were used to define their boundaries. Six thought categories were analyzed, excluding fillers: current state, semantic-world, semantic-self, episodic, future-oriented, and other (Fig. 3a). For each category, we calculated the proportion of each of the six categories immediately following it. This resulted in a six-by-six transition probability matrix for each participant, where each row represents the current category and each column represents the next category. If a participant’s response did not contain a particular category, the transition probabilities from that category (i.e., the row for that category) were considered nonexistent and excluded from the analysis. We then tested whether specific transitions between categories occurred more frequently than expected by chance. For each pair of current and next categories in the transition probability matrix, we performed a two-tailed paired-samples *t*-test, comparing the transition probabilities to the overall proportion of the next category among all sentences generated within each participant.

### Semantic similarity between thoughts

To quantify semantic similarity between thoughts, we employed a natural language processing technique that transforms text into embedding vectors. We used a pretrained model (all-mpnet-base-v2) implemented in the Sentence Transformers Python module (version 2.2.0; https://www.sbert.net) to convert the transcribed text of each thought unit into a 768-dimensional vector. Semantic similarity between pairs of thought units was then defined as the cosine similarity between their respective embedding vectors (Fig. 3b).

To examine the effect of temporal proximity on semantic similarity between thoughts, we computed the semantic similarity between each thought unit (i.e., target) and the 15 thoughts preceding and following the target within each participant (Fig. 3c). The semantic similarity as a function of lag from the target was averaged across all target thoughts within each of the five thought categories, excluding “filler” and “other”. To directly compare thoughts that are near and far from the target, we averaged the semantic similarity at lags 1 and −1 (“near”) and at lags 15 and −15 (“far”) within each participant and thought category. We then performed two-tailed paired-samples *t*-tests for each category, using lag (near, far) as a within-participant factor.

To compare the semantic similarity at different types of thought boundaries, we averaged the semantic similarity between consecutive thoughts within each type of boundary: 1) where only the category of thoughts changed, 2) where only the topic of thoughts changed, and 3) where both the category and topic changed. The averaging was done for each participant. We then performed a one-way repeated-measures ANOVA with thought boundary type as a within-participant factor.

### Thought boundary agreement

To measure the strength of boundaries between thoughts without explicitly considering thought categories or topics, a separate group of human coders manually identified boundaries within the 118 fMRI participants’ think-aloud responses. We recruited 185 coders online from Sona and Prolific, compensating them with course credits or monetary rewards. An additional 19 coders were excluded due to failure to follow instructions. Each coder read an average of 3.83 think-aloud transcripts (range: 1 – 10), with each row corresponding to a sentence or clause containing a single thought category and topic. The coders were instructed to use their best subjective judgement to segment each transcript into individual thought units by assigning different numbers to rows representing different thoughts (Fig. 3d). The offset of the last sentence/clause of one thought before a new thought began was identified as the thought boundary. The coders also identified filler utterances that did not correspond to specific thoughts; changes to or from fillers were not considered as thought boundaries. Each transcript was segmented by an average of 6 coders (range: 5 – 8), and the proportion of coders who identified a moment as a boundary (i.e., boundary agreement) served as a measure of boundary strength.

To compare boundary strength at different types of thought boundaries (i.e., category change only, topic change only, both change) as defined by the manual category and topic coding, we averaged the boundary agreement scores within each participant for each boundary type. We then performed a one-way repeated-measures ANOVA with thought boundary type as a within-participant factor.

### MRI data acquisition

MRI scanning was conducted at the F. M. Kirby Research Center for Functional Brain Imaging at Kennedy Krieger Institute on a 3 Tesla Philips Ingenia Elition scanner with a 32-channel head coil. Functional images were acquired using a T2*-weighted multiband accelerated echo-planar imaging (EPI) sequence (TR = 1.5 s; TE = 30 ms; flip angle = 52°; acceleration factor = 4; 60 oblique axial slices; grid size 112 × 112; voxel size 2 × 2 × 2 mm^3^). Fieldmap images were also acquired to correct for B0 magnetic field inhomogeneity (60 oblique axial slices; grid size 112 × 112; voxel size 2 × 2 × 2 mm^3^). Whole-brain high-resolution anatomical images were acquired using a T1-weighted MPRAGE pulse sequence (150 axial slices; grid size 224 × 224; voxel size 1 × 1 × 1 mm^3^).

### MRI data preprocessing

MRI data collected during think-aloud sessions were first organized into the Brain Imaging Data Structure (BIDS) format using custom scripts. Preprocessing of high-resolution anatomical images and cortical surface reconstruction were performed using the recon-all pipeline of FreeSurfer^74^. Functional images were preprocessed using fMRIprep^75^ (version 21.0.2; RRID:SCR_016216) with default settings. Specifically, functional images were corrected for head motion and B0 magnetic inhomogeneity. Functional images were then coregistered to the anatomical image and resampled to the fsaverage6 template surface (for cortical analysis) and the MNI 152 volume space (for subcortical analysis). Additionally, functional images were smoothed (FWHM 4 mm) in the surface space using FreeSurfer’s mri_surf2surf and in the volume space using FSL’s SUSAN^76^ (Smoothing over Univalue Segment Assimilating Nucleus;). Nuisance regressors including linear and quadratic trends, six head motion parameters (translation and rotation in the x, y, and z dimensions), and the average time courses of the whole-brain mask, cerebrospinal fluid, and white matter were then projected out from the smoothed data. The resulting time series were z-scored within each vertex or voxel across all volumes within the scanning run. Within each scanning run, the first 10 volumes were discarded. Motion outlier volumes (framewise displacement >= 1 mm), along with the two volumes immediately preceding and following each outlier, were also excluded from the analysis.

### Cortical parcellation and region of interest (ROI) definition

For whole-brain activation and pattern similarity analyses, we used a cortical parcellation atlas based on functional connectivity patterns identified through fMRI^35^. The atlas divides the cortical surface into 400 parcels, with 200 parcels in each hemisphere, which are grouped into 17 functional networks identified in a previous study^76^.

For region-of-interest analyses, we defined the bilateral posterior-medial cortex (PMC) and the bilateral auditory cortex (Fig. 4a) by combining parcels from the 400-parcel atlas that correspond to the regions. The PMC ROI included parcels from the posterior cingulate cortex and precuneus within Default Network A. The auditory cortex ROI consisted of parcels around the primary and secondary auditory cortices within Somatomotor Network B (see Supplementary Table 8 for the list of parcels included in the ROIs). The bilateral hippocampus mask was obtained from the subcortical atlas (Aseg) provided by FreeSurfer, using the MNI volume space as reference.

For functional connectivity analysis, we extracted individual subregions from three a priori functional networks of interest out of the 17 networks in the atlas: Control Network B, Default Network A, and Default Network C (Fig. 5d). Parcels corresponding to each subregion defined by the atlas were combined to form a single region. Control Network B consisted of subregions in the interior parietal lobule (IPL), dorsal prefrontal cortex (PFCd), lateral dorsal prefrontal cortex (PFCld), lateral ventral prefrontal cortex (PFClv), medial posterior prefrontal cortex (PFCmp), and temporal lobe. Default Network A consisted of subregions in the IPL, PFCd, medial prefrontal cortex (PFCm), temporal lobe, and PMC. Default Network C consisted subregions in the IPL, parahippocampal cortex (PHC), and retrosplenial cortex (Rsp). Additionally, we included the hippocampus as a subregion of Default Network C.

### Univariate activation for different thought categories

We performed whole-brain univariate activation analysis to identify regions recruited during spontaneous memory recall and future thinking. For each participant, we computed the mean activation for each thought category within each cortical parcel from the 400-parcel atlas. This was done by first averaging the preprocessed blood oxygenation level-dependent (BOLD) signal across all vertices within a parcel and across TRs within each thought unit. We then averaged these mean signals across thought units corresponding to each thought category. Next, for each parcel, we performed group-level contrasts between the “current state” category and each of the other thought categories of interest (i.e., semantic-world, semantic-self, episodic, and future-oriented) using two-tailed paired-samples *t*-tests. This resulted in whole-brain *t*-statistic and *p*-statistic maps for each of the four contrasts. We applied Bonferroni’s correction to each contrast map to account for multiple comparisons across all 400 parcels.

We additionally compared univariate activation across thought categories within the PMC and hippocampus ROIs. For each participant and ROI, we computed the mean activation for each thought category by averaging the preprocessed BOLD signal across vertices/voxels and across TRs within each thought unit, and then averaging across all thought units corresponding to each category. For each ROI, we performed a one-way repeated-measures ANOVA with the thought category as a within-subject factor to test for statistically significant differences in activation across thought categories. For both whole-brain and ROI analyses, the time windows corresponding to individual thought units were shifted forward by 4.5 seconds to account for the hemodynamic response delay.

### Univariate activation at thought boundaries

We first performed a whole-brain analysis (Fig. 4a) to identify brain regions activated at strong thought boundaries, defined as those with boundary agreement scores of 1. For each cortical parcel in each participant, we computed the mean activation during strong thought boundary periods and non-boundary periods. A strong boundary period was defined as a 6-second window starting at the offset of the thought that immediately preceded a strong boundary. A non-boundary period was defined as the 6-second window in the middle of thoughts lasting longer than 15 seconds. To account for the hemodynamic response delay, both boundary and non-boundary time windows were shifted forward by 4.5 seconds. Preprocessed BOLD signals were first averaged across all TRs within the boundary/non-boundary periods and then across vertices within each parcel. Next, for each parcel, we performed a group-level contrast between the strong boundary periods and non-boundary periods using two-tailed paired-samples *t*-tests. The resulting whole-brain *t*-statistic map was corrected for multiple comparisons across all 400 parcels using Bonferroni’s method.

We also examined activation time courses evoked by different types of thought boundaries in the PMC and hippocampus ROIs. For each participant and ROI, we averaged TR-by-TR activation across all vertices/voxels within the ROI. From this activation time series, we extracted 27-second (18 TRs) time courses locked to thought boundaries (i.e., from 2 TRs before to 15 TRs after the thought offset TR). The time courses were then averaged across boundaries within each boundary type: 1) strong thought boundaries with boundary agreement scores of 1, 2) boundaries where only the topic of thoughts changed, and 3) boundaries where only the category of thoughts changed. For the non-boundary control condition, we additionally extracted and averaged time courses locked to the middle of thoughts lasting longer than 15 seconds (i.e., from 2 TRs before to 15 TRs after the middle TR). Two-tailed paired-samples *t*-test were performed to compare activation levels between boundary and non-boundary conditions at each time point of the time courses. Bonferroni’s correction was applied to correct for multiple comparisons across the 18 time points.

### Distributed activation pattern at thought boundaries

We conducted a whole-brain pattern similarity analysis to test if the major mental context transition pattern observed in our prior study^7^ generalized to strong thought boundaries during the think-aloud task (Fig. 4c). We first extracted strong thought boundary and non-boundary patterns for each cortical parcel in each participant’s brain. A strong thought boundary pattern was generated by averaging activation patterns across all TRs within 6-second windows starting from the offset of thoughts immediately preceding strong boundaries with an agreement score of 1. A non-boundary pattern was generated by averaging activation patterns across all TRs within the middle 6 seconds of thoughts lasting longer than 15 seconds. To account for the hemodynamic response delay, both boundary and non-boundary time windows were shifted forward by 4.5 seconds.

We next computed Pearson correlations between the think-aloud boundary/non-boundary patterns and the between-movie boundary pattern obtained from the movie watching phase data of our prior fMRI study^77^. In that study, participant watched a series of ten short (2 – 8 minutes long) audiovisual movie stimuli, separated by 6-second title scenes. Participants subsequently verbally recalled the movies in any order they wanted. We followed the procedures described in ref.^7^ to generate the between-movie boundary pattern for each participant. Activation patterns were first averaged across time points within the 15-second boundary period following the offset of each movie, shifted forward by 4.5 seconds. The patterns were then averaged across movie stimuli and fifteen participants analyzed in the dataset to create a single template pattern. To preserve the boundary activation pattern reported in the original study as much as possible, we used the dataset preprocessed according to the pipeline described in ref.^7^.

To test whether the strong boundary patterns were overall positively correlated with the between-movie boundary patterns, we performed group-level two-tailed one-sample *t*-tests against zero on the correlation coefficients for each cortical parcel. Additionally, group-level two-tailed paired-samples *t*-tests were performed to directly compare the similarity of the between-movie boundary patterns to the strong thought boundary patterns versus the non-boundary patterns. Bonferroni’s correction was applied to each resulting whole-brain statistical parametric map to correct for multiple comparisons across parcels. Finally, to identify parcels showing significant effects in both tests after correction, we masked the areas with higher similarity to the strong boundary pattern compared to the non-boundary pattern with the areas that showed overall positive similarity to the strong boundary pattern. We also performed the same boundary pattern similarity analysis within the PMC and auditory cortex ROIs, as was done for individual parcels in the whole-brain analysis.

Finally, we tested whether the strong thought boundary patterns in the PMC and auditory ROIs were influenced by temporary silence due to pauses in speech at the boundaries. To do this, we compared the think-aloud boundary/non-boundary patterns with the activation pattern associated with silence, measured during the movie watching phase of ref.^7^. Silent periods were identified as moments within movies when the audio amplitude, convolved with a hemodynamic response function, was as low as the mean amplitude during the title scenes between movies where no sound was presented. To prevent potential carryover effects from the between-movie boundaries, time points within the first 45 seconds of each movie were excluded from the silent periods. Activation patterns were averaged across all time points within the silent periods for each participant and then across participants. We performed group-level two-tailed one-sample *t*-tests against zero to test the overall positivity of correlations between the silence template pattern and the think-aloud boundary/non-boundary patterns. Additionally, we performed two-tailed paired-samples *t*-tests to compare the similarity of the silence template pattern to the strong thought boundary versus the non-boundary patterns.

### Thought network structure

To quantify the overall structure of think-aloud responses, we transformed each participant’s response into a semantic network (Fig. 5a) following the procedures developed in our prior study^77^. In this network, the nodes represented individual thought units, and the edges between nodes represented the semantic similarity between thoughts. Semantic similarity between all possible pairs of thoughts was measured using the procedures described above in the “Semantic similarity between thoughts” section. Specifically, each thought unit was converted into a 768-dimensional text embedding vector, and the cosine similarity between these vectors was computed. The resulting network was undirected, and edges with weights below zero were removed. To measure the global structure of the thought network, we calculated the average clustering coefficient across all nodes (Fig. 5b). The clustering coefficient for each node was defined as the geometric average of the subgraph edge weights, using the implementation provided by the NetworkX Python package (version 3.1; https://networkx.org/).

### Functional connectivity analysis

To examine how interactions between brain regions influence the overall structure of thought networks, we computed functional connectivity within and between brain networks involved in spontaneous thought generation^10,23,52^. Specifically, we focused on Control Network B, Default Network A, and Default Network C as defined in the 17-network version of the 400-parcel cortical atlas^35^. These networks anatomically overlap with the frontoparietal control network, the core default network subsystem, and the medial temporal lobe subsystem of the default network discussed in prior studies^10,52^, respectively.

Functional connectivity was computed from the entire think-aloud session for each participant. First, we extracted the mean activation time course of each network subregion by averaging across all vertices/voxels within each region and hemisphere. For bilateral subregions, time courses were also averaged across hemispheres (see Fig. 5d and the “Cortical parcellation and region of interest definition” section above for the list of subregions). Next, we computed pairwise Pearson correlations between the activation time courses of individual subregions. Within-network functional connectivity was then defined as the average of correlations between different subregions within the same network. Between-network functional connectivity was defined as the average of correlations between all possible pairs of subregions across two different networks.

Finally, we computed Pearson correlations between the participant-wise average thought network clustering coefficients and the within/between-network functional connectivity values. As a post-hoc exploration, we also computed correlations between the thought network clustering coefficients and the functional connectivity between all individual subregions in the three brain networks. Bonferroni’s correction was applied to correct for multiple comparisons across all possible pairs of subregions.

### Statistical tests

Details of the statistical tests used in each analysis are provided in the corresponding subsection of the Methods section. All statistical tests were two-tailed. For parametric tests comparing means across multiple conditions, Mauchly’s test was used to assess sphericity. If the assumption of sphericity was violated, the Greenhouse-Geisser corrected *p*-value was reported in place of the uncorrected *p*-value. Only participants with complete data for all relevant conditions were included in comparisons. Participants with missing data in any condition (e.g., those who did not generate thoughts in the current state category when comparing current state and episodic recall) were excluded from the relevant comparisons.

## DATA AVAILABILITY

The raw fMRI and behavioral data from think-aloud sessions will be made publicly available on OpenNeuro.org following the publication of this study.

## CODE AVAILABILITY

The analyses in this study were conducted using custom Python scripts and publicly available packages. These analysis scripts will be made available upon request to the corresponding author (H. L.).

## ACKNOWLEDGEMENTS

We thank Yoonjung Lee for assisting with collecting fMRI data and Colette Youstra for assisting with collecting behavioral thought segmentation data. C. J. H. was supported by National Institute of Mental Health (R01MH119099) and National Science Foundation CAREER Award (BCS-2238711). J. C. was supported by National Institute of Mental Health (R01MH133732).

