## Supplementary Materials for "Neural dynamics of spontaneous memory recall and future thinking in the continuous flow of thoughts"

### SUPPLEMENTARY TABLES

**Supplementary Table 1.** Mean duration, word count, speech rate (number of words/second) per thought unit, and streak length (number of consecutive thought units) for each thought category.

| Thought category | Number of participants | Duration in seconds | Word count | Speech rate | Streak length |
|---|---|---|---|---|---|
| Current | 98 | 8.96 (6.78) | 21.85 (15.44) | 2.88 (.74) | 1.3 (.45) |
| Semantic (world) | 117 | 11.7 (6.59) | 29.83 (15.44) | 2.98 (.6) | 1.25 (.26) |
| Semantic (self) | 113 | 7.48 (4.1) | 21.04 (10.82) | 3.34 (.77) | 1.16 (.23) |
| Episodic | 113 | 10.89 (6.12) | 29.07 (15.73) | 3.07 (.74) | 1.25 (.4) |
| Future | 112 | 9.45 (4.42) | 24.75 (11.33) | 3.13 (.92) | 1.67 (1.88) |
| Other | 20 | 4.48 (3.36) | 12.13 (6.36) | 3.84 (1.73) | 1.01 (.03) |

Numbers in parentheses represent standard deviations across participants.

**Supplementary Table 2.** List of suprathreshold parcels from the univariate contrast between Semantic memory about the world/other people and Current state.

| Direction of effect | Hemisphere | Network | Parcel | *t* * | *p* (unc.) |
|---|---|---|---|---|---|
| Semantic (world) > Current state | Left | DefaultA | IPL_1 | 4.46 | .00003636 |
| | | | pCunPCC_1 | 5.3 | .00000167 |
| | | SomMotB | Cent_3 | 4.27 | .00007046 |
| | Right | DefaultA | pCunPCC_1 | 4.59 | .00002285 |
| | | SomMotA | SomMotA_10 | 5.87 | .00000019 |
| Current state > Semantic (world) | Left | ContA | IPS_2 | -5.57 | .00000061 |
| | | ContC | pCun_1 | -5.85 | .00000021 |
| | | | pCun_2 | -4.81 | .00001026 |
| | | DorsAttnB | PostC_7 | -4.74 | .00001343 |
| | | SalVentAttnA | ParOper_3 | -5.33 | .00000149 |
| | Right | ContB | IPL_1 | -5.34 | .00000147 |
| | | ContC | pCun_2 | -5.12 | .00000327 |
| | | DorsAttnA | SPL_3 | -5.13 | .00000319 |
| | | SalVentAttnB | IPL_1 | -6.09 | .00000008 |
| | | | PFClv_1 | -4.44 | .00003873 |
| | | TempPar | TempPar_10 | -4.9 | .00000739 |

*Degree of freedom = 61

**Supplementary Table 3.** List of suprathreshold parcels from the univariate contrast between Semantic memory about oneself and Current state.

| Direction of effect | Hemisphere | Network | Parcel | *t* * | *p* (unc.) |
|---|---|---|---|---|---|
| Semantic (self) > Current state | Left | DefaultA | IPL_1 | 5.1 | .00000389 |
| | | | pCunPCC_1 | 5.19 | .00000277 |
| | | DefaultB | Temp_3 | 4.55 | .00002828 |
| | | SomMotB | Cent_3 | 4.34 | .00005838 |
| | Right | SomMotA | SomMotA_10 | 4.73 | .00001482 |
| | | SomMotB | S2_4 | 4.44 | .0000403 |
| Current state > Semantic (self) | Left | ContA | IPS_2 | -4.97 | .0000062 |
| | | ContB | IPL_3 | -4.76 | .00001332 |
| | | ContC | pCun_1 | -5.97 | .00000015 |
| | | | pCun_2 | -5.78 | .00000031 |
| | | | pCun_3 | -5.44 | .00000111 |
| | | SalVentAttnA | ParOper_3 | -5.13 | .0000035 |
| | Right | ContB | IPL_1 | -4.19 | .00009479 |
| | | ContC | pCun_2 | -5.73 | .00000038 |
| | | | pCun_5 | -4.73 | .00001461 |
| | | DorsAttnA | SPL_3 | -4.73 | .0000149 |
| | | SalVentAttnB | IPL_1 | -4.42 | .00004335 |

*Degree of freedom = 58

**Supplementary Table 4.** List of suprathreshold parcels from the univariate contrast between Episodic recall and Current state.

| Direction of effect | Hemisphere | Network | Parcel | *t* * | *p* (unc.) |
|---|---|---|---|---|---|
| Episodic > Current state | Left | DefaultA | IPL_1 | 5.42 | .00000114 |
| | | | IPL_2 | 6.97 | < .00000001 |
| | | | pCunPCC_1 | 7.61 | < .00000001 |
| | | | pCunPCC_2 | 4.38 | .00004925 |
| | | DefaultC | Rsp_1 | 4.81 | .00001078 |
| | | | Rsp_2 | 6.26 | .00000005 |
| | Right | DefaultA | IPL_2 | 5.44 | .0000011 |
| | | | PFCd_2 | 4.83 | .00001004 |
| | | | pCunPCC_1 | 6.78 | .00000001 |
| | | SomMotA | SomMotA_10 | 4.37 | .00005138 |
| Current state > Episodic | Left | ContA | IPS_2 | -7.06 | < .00000001 |
| | | ContC | pCun_2 | -4.69 | .0000165 |
| | | DorsAttnA | SPL_6 | -4.25 | .00007702 |
| | | SalVentAttnA | ParOper_3 | -5.11 | .00000371 |
| | Right | ContB | IPL_1 | -4.24 | .00008098 |
| | | ContC | pCun_2 | -4.15 | .00010936 |
| | | | pCun_5 | -4.11 | .0001244 |
| | | DorsAttnA | SPL_3 | -5.73 | .00000036 |
| | | SalVentAttnB | IPL_1 | -5.38 | .00000136 |
| | | TempPar | TempPar_10 | -4.37 | .00005083 |
| | | VisCent | ExStr_11 | -4.44 | .00004019 |
| | | | ExStr_9 | -4.96 | .00000633 |
| | | VisPeri | ExStrInf_1 | -4.42 | .00004327 |

*Degree of freedom = 59

**Supplementary Table 5.** List of suprathreshold parcels from the univariate contrast between Future-oriented thinking and Current state.

| Direction of effect | Hemisphere | Network | Parcel | *t* * | *p* (unc.) |
|---|---|---|---|---|---|
| Future thinking > Current state | Left | ContB | PFCd_1 | 5 | .00000543 |
| | | DefaultA | IPL_1 | 5.86 | .00000022 |
| | | | IPL_2 | 7.12 | < .00000001 |
| | | | PFCd_2 | 6.3 | .00000004 |
| | | | PFCd_3 | 5.26 | .00000208 |
| | | | PFCm_1 | 5.91 | .00000018 |
| | | | PFCm_2 | 6.66 | .00000001 |
| | | | pCunPCC_1 | 10.28 | < .00000001 |
| | | | pCunPCC_2 | 7.93 | < .00000001 |
| | | | pCunPCC_3 | 6.1 | .00000009 |
| | | DefaultC | Rsp_1 | 6.01 | .00000013 |
| | | | Rsp_2 | 7.87 | < .00000001 |
| | Right | ContB | PFCld_4 | 4.8 | .00001113 |
| | | DefaultA | IPL_2 | 4.89 | .00000817 |
| | | | PFCd_2 | 7.2 | < .00000001 |
| | | | PFCm_1 | 5.25 | .00000218 |
| | | | pCunPCC_1 | 9.47 | < .00000001 |
| | | | pCunPCC_2 | 6.02 | .00000012 |
| | | | pCunPCC_3 | 5.69 | .00000042 |
| | | DefaultC | Rsp_2 | 4.16 | .00010486 |
| | | SomMotA | SomMotA_10 | 4.88 | .00000837 |
| Current state > Future thinking | Left | ContA | IPS_2 | -5.32 | .00000167 |
| | | DefaultB | PFCv_5 | -4.29 | .00006669 |
| | | SalVentAttnA | FrOper_2 | -4.61 | .0000218 |
| | | | ParOper_3 | -7.87 | < .00000001 |

| | | | | | |
|---|---|---|---|---|---|
| | Right | SalVentAttnB | IPL_1 | -4.66 | .00001834 |
| | | | PFClv_1 | -5.25 | .00000219 |

*Degree of freedom = 59

**Supplementary Table 6.** Mean percentages of pre- and post-boundary thought category pairs at strong thought boundaries with a boundary agreement score of 1.

| | | Post-boundary category | | | | | |
|---|---|---|---|---|---|---|---|
| | | Current | Semantic (world) | Semantic (self) | Episodic | Future | Other |
| Pre-boundary category | Current | 3.49 (10.30) | 2.51 (7.12) | 1.42 (4.44) | 3.54 (10.79) | 3.61 (7.77) | .05 (.51) |
| | Semantic (world) | 4.91 (8.49) | 6.29 (10.32) | 3.70 (8.20) | 4.90 (12.75) | 7.59 (13.64) | .48 (2.37) |
| | Semantic (self) | 4.06 (10.11) | 4.86 (12.61) | 3.10 (8.62) | 2.19 (5.77) | 3.96 (7.95) | .22 (1.39) |
| | Episodic | 2.27 (6.93) | 2.55 (10.02) | 1.65 (5.30) | 2.40 (6.82) | 1.77 (4.43) | 0 (0) |
| | Future | 2.80 (6.36) | 5.99 (10.67) | 1.53 (4.32) | 1.96 (5.13) | 14.90 (20.74) | .18 (1.37) |
| | Other | .18 (1.37) | .13 (1.05) | 0 (0) | .05 (.51) | .05 (.51) | 0 (0) |

N = 117, excluding one participant whose think-aloud response did not include a strong boundary. Numbers in parentheses represent standard deviations across participants.

**Supplementary Table 7.** List of suprathreshold parcels from the univariate contrast between Strong boundary periods and Non-boundary periods.

| Direction of effect | Hemisphere | Network | Parcel | *t* * | *p* (unc.) |
|---|---|---|---|---|---|
| Strong boundary > Non-boundary | Left | ContC | Cingp_1 | 7.75 | < .00000001 |
| | | | Cingp_2 | 6.5 | .00000001 |
| | | | pCun_1 | 8.33 | < .00000001 |
| | | | pCun_2 | 4.34 | .00004528 |
| | | DefaultA | PFCm_4 | 5.14 | .0000022 |
| | | | PFCm_6 | 4.79 | .0000083 |
| | | | pCunPCC_1 | 5.66 | .00000028 |
| | | | pCunPCC_2 | 5.4 | .00000078 |
| | | | pCunPCC_3 | 6.42 | .00000001 |
| | | | pCunPCC_4 | 4.9 | .0000055 |
| | | | pCunPCC_6 | 4.3 | .00005187 |
| | | DefaultC | Rsp_1 | 4.82 | .0000076 |
| | | | Rsp_2 | 4.6 | .00001721 |
| | Right | ContC | Cingp_1 | 6.74 | < .00000001 |
| | | | Cingp_2 | 6.43 | .00000001 |
| | | | pCun_2 | 5.49 | .00000055 |
| | | DefaultA | pCunPCC_1 | 4.45 | .00002931 |
| | | | pCunPCC_5 | 5.14 | .00000219 |
| Non-boundary > Strong boundary | Left | DorsAttnA | TempOcc_1 | -4.46 | .00002863 |
| | | | TempOcc_3 | -4.57 | .000019 |
| | | DorsAttnB | PrCv_1 | -5.03 | .00000334 |
| | | SalVentAttnA | FrOper_2 | -4.7 | .00001169 |
| | | SomMotA | SomMotA_17 | -4.07 | .00011507 |
| | | SomMotB | Aud_1 | -4.28 | .00005622 |
| | | | Aud_2 | -5.05 | .00000313 |

| | | | | | |
|---|---|---|---|---|---|
| | | | Aud_3 | -5.29 | .00000118 |
| | | | Aud_4 | -5.15 | .00000208 |
| | | | Cent_1 | -4.51 | .00002348 |
| | | | Cent_2 | -4.34 | .00004442 |
| | | | Cent_3 | -4.72 | .00001104 |
| | | | Cent_4 | -6.9 | < .00000001 |
| | | | Cent_5 | -5.98 | .00000007 |
| | | | S2_3 | -4.19 | .0000759 |
| | | TempPar | TempPar_2 | -5.89 | .00000011 |
| | | | TempPar_3 | -5.89 | .00000011 |
| | | | TempPar_4 | -5.43 | .00000069 |
| | | | TempPar_5 | -5.02 | .00000348 |
| | Right | DorsAttnA | TempOcc_2 | -4.33 | .00004641 |
| | | SomMotA | SomMotA_14 | -4.37 | .0000397 |
| | | | SomMotA_19 | -4.91 | .0000052 |
| | | SomMotB | Aud_2 | -5.08 | .00000278 |
| | | | Aud_3 | -6.39 | .00000001 |
| | | | Cent_2 | -4.25 | .00006163 |
| | | | Cent_3 | -5.01 | .00000363 |
| | | | S2_4 | -4.56 | .00001975 |
| | | TempPar | TempPar_4 | -4.15 | .00008896 |

*Degree of freedom = 74

**Supplementary Table 8.** List of Schaefer 400 parcels used to create regions of interest.

| Region of interest | Hemisphere | Network | Parcel |
|---|---|---|---|
| Posterior medial cortex | Left | DefaultA | pCunPCC_1 |
| | | | pCunPCC_2 |
| | | | pCunPCC_3 |
| | | | pCunPCC_4 |
| | | | pCunPCC_5 |
| | | | pCunPCC_6 |
| | | | pCunPCC_7 |
| | Right | DefaultA | pCunPCC_1 |
| | | | pCunPCC_2 |
| | | | pCunPCC_3 |
| | | | pCunPCC_4 |
| | | | pCunPCC_5 |
| Auditory cortex | Left | SomMotB | Aud_1 |
| | | | Aud_2 |
| | | | Aud_3 |
| | | | Aud_4 |
| | | | Ins_1 |
| | | | S2_2 |
| | | | S2_5 |
| | Right | SomMotB | Aud_1 |
| | | | Aud_2 |
| | | | Aud_3 |
| | | | Ins_1 |
| | | | S2_2 |
| | | | S2_4 |

### SUPPLEMENTARY FIGURES

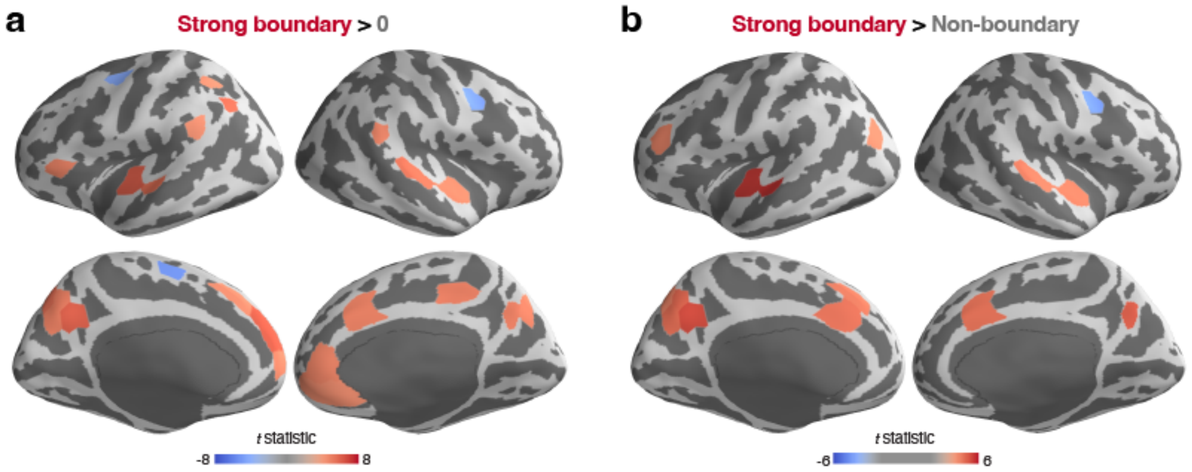


**Suppl. Fig. 1. Whole-brain pattern similarity with the between-movie boundary pattern. a.** Whole-brain *t*-statistic map of the pattern similarity between the strong thought boundary pattern during think-aloud and the between-movie boundary pattern during movie watching[1], compared against zero. Parcels with correlation coefficients significantly greater than zero are shown in red, while those with coefficients significantly smaller than zero are shown in blue. **b.** Whole-brain *t*-statistic map of the pattern similarity contrast between the strong thought boundary period and the non-boundary period. Parcels whose between-movie boundary patterns are significantly more similar to their strong thought boundary patterns than to their non-boundary patterns are shown in red, while those that are less similar are shown in blue. In both **a** and **b**, The *t*-statistic maps are displayed on the lateral (top) and medial (bottom) surfaces of the inflated fsaverage6 template brain. Statistical significance ($p < .05$) was Bonferroni corrected across all parcels.

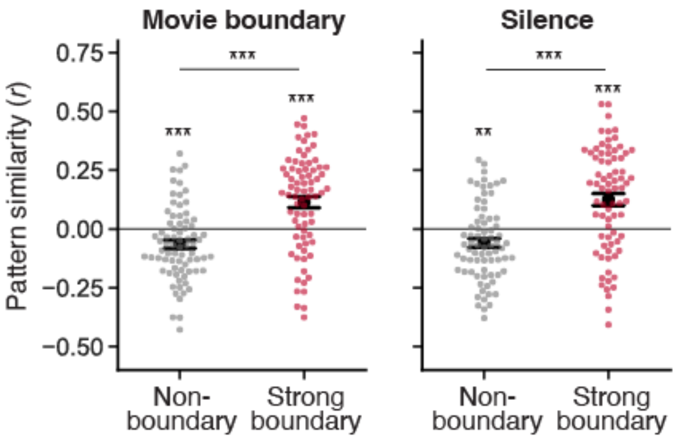


**Suppl. Fig. 2. Boundary pattern similarity in the auditory cortex.** The think-aloud strong thought boundary and non-boundary patterns were correlated with the mean activation patterns of between-movie boundary periods (left panel) or silent periods (right panel) from the movie watching phase[1]. Each colored dot represents an individual participant (N = 75). Black circles indicate the mean across participants. Error bars show the SEM across participants. $^{**}p < .01$, $^{***}p < .001$ (uncorrected).
